# The contribution of Neanderthal introgression and natural selection to neurodegenerative diseases

**DOI:** 10.1101/2022.04.29.490053

**Authors:** Zhongbo Chen, Regina H. Reynolds, Antonio F. Pardiñas, Sarah A. Gagliano Taliun, Wouter van Rheenen, Kuang Lin, Aleksey Shatunov, Emil K. Gustavsson, Isabella Fogh, Ashley R. Jones, Wim Robberecht, Philippe Corcia, Adriano Chiò, Pamela J. Shaw, Karen E. Morrison, Jan H. Veldink, Leonard H. van den Berg, Christopher E. Shaw, John F. Powell, Vincenzo Silani, John A. Hardy, Henry Houlden, Michael J. Owen, Martin R. Turner, Mina Ryten, Ammar Al-Chalabi

## Abstract

Humans are thought to be more susceptible to neurodegeneration than equivalently-aged primates. It is not known whether this vulnerability is specific to anatomically-modern humans or shared with other hominids. The contribution of introgressed Neanderthal DNA to neurodegenerative disorders remains uncertain. It is also unclear how common variants associated with neurodegenerative disease risk are maintained by natural selection in the population despite their deleterious effects. In this study, we aimed to quantify the genome-wide contribution of Neanderthal introgression and positive selection to the heritability of complex neurodegenerative disorders to address these questions.

We used stratified-linkage disequilibrium score regression to investigate the relationship between five SNP-based signatures of natural selection, reflecting different timepoints of evolution, and genome-wide associated variants of the three most prevalent neurodegenerative disorders: Alzheimer’s disease, amyotrophic lateral sclerosis and Parkinson’s disease.

We found no evidence for enrichment of positively-selected SNPs in the heritability of Alzheimer’s disease, amyotrophic lateral sclerosis and Parkinson’s disease,, suggesting that common deleterious disease variants are unlikely to be maintained by positive selection. There was no enrichment of Neanderthal introgression in the SNP-heritability of these disorders, suggesting that Neanderthal admixture is unlikely to have contributed to disease risk.

These findings provide insight into the origins of neurodegenerative disorders within the evolution of *Homo sapiens* and addresses a long-standing debate, showing that Neanderthal admixture is unlikely to have contributed to common genetic risk of neurodegeneration in anatomically-modern humans.

## Introduction

Encephalisation and the evolution of complex human-specific traits are thought to have increased the susceptibility of *Homo sapiens* to disorders of the brain compared to their aged non-human primate counterparts. ^1–3^ This is seen in Alzheimer’s and Parkinson’s disease, which seldom occur naturally on a pathological or phenotypic level in non-human species.^1,2^ Likewise, unique motor dysfunction in amyotrophic lateral sclerosis (ALS) supports the selective vulnerability of the highly-developed corticomotoneuronal system in humans.^4–6^ Human-lineage-specific genomic sequences have been shown to be enriched for brain-specific elements and risk loci for neurodegenerative disorders.^7^ Thus, while positive natural selection may have driven a proportion of human-specific adaptive evolution,^8,9^ it may be possible that the same selected variants also influence the risk of neurodegenerative disease.

It is not known whether this neuro-vulnerability arose after divergence from other species or whether it represents a more recent phenomenon, characteristic of modern-day humans over other hominids. As anatomically-modern humans migrated out of Africa 50 to 100 thousand years ago, they interbred with archaic hominins including Neanderthals.^10^ As a result, Neanderthal DNA accounts for approximately 1–4% of the modern Eurasian genome.^11–13^ While most Neanderthal DNA experienced purifying selective pressures,^14^ positive selection of these archaic alleles may have contributed to modern human adaptation to the non-African environment^15^ through modulating dermatological^14^, immunological^16,17^ and metabolic function.^18,19^ These introgressed Neanderthal alleles have also been implicated in contributing to the risk of some conditions, including actinic keratosis, depression and obesity.^20^

It remains unclear how much Neanderthal admixture has affected our risk of neurodegenerative disorders. While Neanderthal single nucleotide polymorphisms (SNPs) may be associated with “neurological” phenotypes in electronic health records of European patients, these nervous system traits were not representative of neurodegenerative diseases.^20^ More recently, two studies aimed to address the complex nature of medically-relevant traits and the genome-wide influence of Neanderthal admixture using UK Biobank data.^21,22^ The former study found that Neanderthal introgression did not significantly contribute to neurological and psychiatric traits and the latter found that introgressed variants were depleted for heritability of high-level cognitive traits.^22^ However, neither looked specifically at neurodegenerative diseases of interest.

Thus, to quantify the contribution of Neanderthal admixture to the heritability of neurodegenerative diseases and to examine whether natural selection maintains common genetic risk of these disorders, we tested the relationship between alleles associated with Alzheimer’s disease^23^, ALS^24^ and Parkinson’s disease^25^ from recent genome-wide association studies (GWAS) with SNP-based signatures of natural selection.^26^ Using stratified-linkage disequilibrium score regression (LDSC), we found that there was no significant enrichment of Neanderthal introgression or positive selection in the SNP-heritability of Alzheimer’s disease, ALS or Parkinson’s disease. Thus, positive selection is unlikely to have played a significant role in the maintenance of common deleterious variants in the genetic architecture of these neurodegenerative diseases.

## Methods

Heritability is defined as the fraction of a trait that is explained by inherited genetic variants in a given environment, and is important for understanding the biology of disease.^27^ More specific to stratified-LDSC, narrow-sense heritability is defined as the proportion of phenotypic variance that can be attributed to variation in the additive effects of genes.^28^ Stratified-LDSC analysis estimates the SNP-based heritability (*h*^2^SNP) of complex traits stratified across different annotations using GWAS data^29,30^ Thus, we were able to use stratified-LDSC to assess the enrichment and depletion of common-*h*^2^SNP of complex neurodegenerative diseases for metrics of Neanderthal introgression and positive selection.^26^

For the metric of Neanderthal introgression, we used the average posterior probability of each human haplotype being the result of Neanderthal admixture estimated by comparing human and Neanderthal genomes (LA).^14,26^ We used four metrics of positive selection: integrated haplotype score (iHS), composite of multiple signals (CMS), cross-population extended haplotype homozygosity (XP-EHH), and composite likelihood ratio (CLR). These metrics were chosen to reflect the different timeframes of the selective processes used and described in previous analyses.^26^ A detailed summary of the calculation of these metrics is shown in **Supplementary Table 1** and code is available in https://github.com/RHReynolds/als-neanderthal-analysis. LA, CLR and CMS were directly retrieved from published references. For both iHS and XP-EHH, we used the metrics calculated previously^26^ using the European superpopulation of the 1,000 Genomes Project Panel 3 dataset, with the African superpopulation used as the second population for XP-EHH. ^26,31^ Further, we transformed the absolute iHS and −log_10_ XP-EHH metrics to ensure that all metrics were on a common scale, in which larger values indicate stronger effect of selection or increased probability of introgression. A summary of the workflow is found in **Figure 1**. iHS measures the amount of extended haplotype homozygosity at a given SNP in the ancestral allele relative to the derived allele and estimates positive selective sweep.^8^ CMS identifies the regions under positive selection by combining long-range haplotypes, differentiated alleles and high frequency derived alleles.^32^ Both iHS and CMS detect more recent selective sweeps in the last 30,000 years.^8,32^ XP-EHH compares two populations to detect an allele that has reached fixation in one population but remains polymorphic in another, identifying alleles that have undergone different selective pressures since population divergence.^33^ Lastly, CLR detects incomplete selective sweeps, quantifying the relative influence of recombination and selection and corrects for background selection.^34^ It can thus detect older signals from ∼ 60,000 to 240,000 years ago.^26^ We did not use any other metrics of purifying selection such as McVicker B-statistic^35^ as a metric of background selection is already incorporated into the baseline LDSC model (v.2.2.).

**Figure 1.**
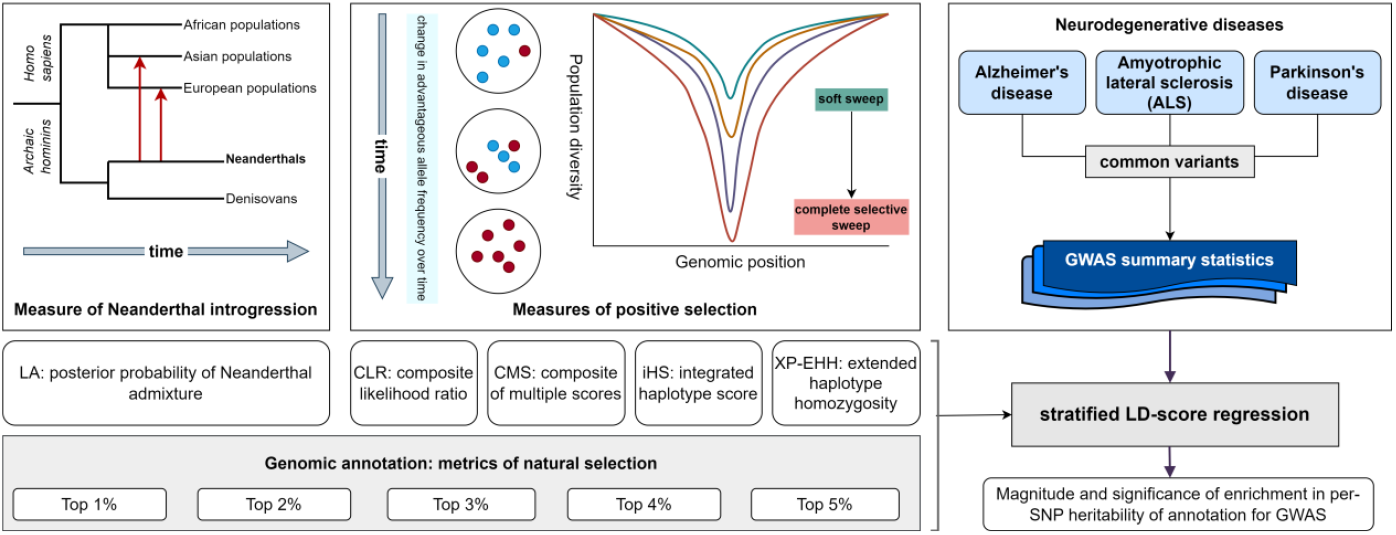
Study workflow. Schematic figure showing key concepts of the study workflow. The red arrows signify admixture from Neanderthals to Eurasian populations; the posterior probability of Neanderthal admixture is measured by LA and partitioned into the top 1–5% metrics and inputted into the stratified-linkage disequilibrium (LD) score regression. Not all gene flow from archaic hominins to *Homo* sapiens is represented in the figure and phylogenetic branches are not to scale. For simplified measures of positive selection, the middle panel shows that with time, an advantageous allele rises in frequency in the population. At the same time, the population diversity decreases at that particular genomic position, and also in other SNPs in linkage disequilibrium that hitchhike with the beneficial allele. This resultant dip in diversity can be shallower in incomplete sweeps and deeper in more complete selective sweeps. Four metrics can measure this positive selection (CLR; CMS; iHS and XP-EHH). These four measures of positive selection were partitioned into the top 1–5% for use in the stratified-LD score regression. We used stratified-LD score regression analysis to test whether these natural selection metrics contribute significantly to heritability of neurodegenerative disease. We assessed the annotation for SNP-heritability contribution in the three most prevalent complex neurodegenerative disorders of Alzheimer’s disease, Parkinson’s disease and amyotrophic lateral scelosis using recent corresponding GWAS summary statistics. Illustrative figures for natural selection are adapted from a review by Vitti *et al*.^39^

We used stratified-LDSC v.1.0.1 (https://github.com/bulik/ldsc/wiki) to test whether these natural selection metrics contribute significantly to heritability of neurodegenerative disease. All natural selection metrics were annotated to the ∼ 9,997,000 SNPs present in the baseline LDSC model (v.2.2), which only includes SNPs with a minor allele frequency of more than 5%. Binary annotations were generated from the natural selection metrics, with thresholds at the top 1%, 2%, 3%, 4% and 5% of the genome-wide values of each metric in the full set of baseline SNPs.^26^ This centile approach was used in previous studies due to difficulties defining the thresholds for selection.^26^ Annotations were then added individually to the baseline LDSC model of 97 annotations (v.2.2, GRCh37),^36^ comprising genome-wide annotations reflecting genetic and LD architecture. HapMap Project Phase 3 (HapMap3)^37^ SNPs and 1000 Genomes Project Phase 3^38^ European population SNPs were used for the regression and LD reference panels, respectively. The major histocompatibility complex region was excluded from all analyses owing to the region’s complicated and long-range LD patterns. This analysis generated a regression coefficient (*τ*_c_), from which we calculated a two-tailed p-value, testing whether the regression coefficient of the annotation category contributes (either through enrichment or depletion) to the trait heritability conditional upon other annotations in the baseline-LD model. False discovery rate (FDR) multiple testing correction was applied across each GWAS, accounting for the number of annotations run (totalling 25 annotations per GWAS). A stringent FDR-corrected p-value (FDR P) < 0.05 was taken to be significant. We assessed the annotation for *h*^*2*^_SNP_ contribution in complex brain-related disorders of apparently sporadic Alzheimer’s disease^23^, Parkinson’s disease (excluding 23&Me participants)^25^ and ALS (European cases only)^24^ using recent corresponding GWAS summary statistics (further description in **Supplementary Table 2**). We chose these three neurodegenerative diseases given that they are the most prevalent of such disorders, have varying mean ages of onset which might affect selection pressures, and have correspondingly well-powered GWAS. All analyses were carried out in R version 4.0.5 (https://www.R-project.org/) and code is available from: https://github.com/RHReynolds/als-neanderthal-analysis.

## Results

To quantify the contribution of Neanderthal introgression and positive natural selection to the heritability of three neurodegenerative disorders, we used stratified-LDSC to assess enrichment or depletion of the h^*2*^SNP of Alzheimer’s disease, ALS and Parkinson’s disease for each SNP-based signature of natural selection. For all analyses, we reported an FDR-corrected two-tailed coefficient p-value that tested whether the regression coefficient (*τ*_c_) of an annotation category contributes to trait heritability, conditional upon other annotations that account for the underlying genetic architecture (**Table 1, Figure 2**, full results in **Supplementary Table 3**).

**Table 1.** Heritability analysis of natural selection metrics. Stratified-LDSC results for SNPs defined by top genome-wide percentiles (1–5%) of all SNPs annotated for each natural selection metric. The regression coefficient (*τ*_c_) represents the contribution of the annotation to trait SNP-heritability (*h*^*2*^_SNP_), controlling for all other annotations within the baseline model. The FDR-corrected p-value (FDR P) is the coefficient p-value following correction for multiple testing. Coefficient FDR P<0.05 and corresponding *τ*_c_ values are highlighted in bold. Neanderthal introgression metric (LA) indicates posterior probability of Neanderthal admixture. Positive selection metrics are composite likelihood ratio statistic (CLR), composite of multiple scores (CMS), integrated haplotype score (iHS), cross-population extended haplotype homozygosity (XP-EHH). Genome-wide association studies (GWAS) include AD2019 (Alzheimer’s disease, Jansen et al. 2019)^23^, ALS2021.EUR (European cases only from amyotrophic lateral sclerosis GWAS, van Rheenen et al. 2021)^24^ and PD2019.met5.ex23&Me (Nalls *et al*. 2019, excluding 23&Me participants).^25^ Full results are shown in **Supplementary Table 3**.

|  |  | Top 1% scores |  | Top 2% scores |  | Top 3% scores |  | Top 4% scores |  | Top 5% scores |  |
| --- | --- | --- | --- | --- | --- | --- | --- | --- | --- | --- | --- |
| GWAS | Metric | $\tau_c$ | FDR P | $\tau_c$ | FDR P | $\tau_c$ | FDR P | $\tau_c$ | FDR P | $\tau_c$ | FDR P |
| Alzheimer's disease (AD2019) | CLR | -1.48E-09 | 0.406 | -2.12E-09 | **0.023** | -2.20E-09 | **0.023** | -2.24E-09 | **0.023** | -1.91E-09 | 0.054 |
|  | CMS | -3.71E-10 | 0.911 | -1.24E-09 | 0.703 | -1.29E-09 | 0.639 | -1.28E-09 | 0.639 | -1.38E-09 | 0.635 |
|  | iHS | -7.29E-09 | 0.520 | -4.17E-09 | 0.520 | -2.16E-09 | 0.635 | -8.97E-10 | 0.753 | -3.72E-10 | 0.908 |
|  | LA | -1.50E-09 | 0.635 | -1.50E-09 | 0.544 | 2.01E-10 | 0.908 | 1.06E-09 | 0.639 | 9.16E-10 | 0.639 |
|  | XP-EHH | -3.58E-09 | 0.563 | -3.97E-09 | 0.406 | -3.26E-09 | 0.406 | -2.90E-09 | 0.406 | -2.21E-09 | 0.518 |
| ALS (ALS2021. EUR) | CLR | -1.11E-08 | 0.160 | -1.01E-08 | 0.160 | -1.01E-08 | 0.160 | -3.76E-09 | 0.852 | -3.75E-09 | 0.852 |
|  | CMS | -5.99E-09 | 0.852 | -4.66E-09 | 0.852 | -3.38E-09 | 0.852 | -1.12E-09 | 0.953 | 3.00E-10 | 0.976 |
|  | iHS | 6.21E-08 | 0.160 | 2.80E-08 | 0.219 | 1.51E-08 | 0.473 | 1.03E-08 | 0.667 | 6.50E-09 | 0.852 |
|  | LA | 2.82E-09 | 0.852 | -1.95E-09 | 0.852 | 2.68E-09 | 0.852 | 2.63E-09 | 0.852 | 1.65E-09 | 0.852 |
|  | XP-EHH | 3.97E-08 | 0.215 | 2.89E-08 | 0.208 | 1.88E-08 | 0.295 | 1.89E-08 | 0.219 | 1.88E-08 | 0.215 |
| Parkinson's disease (PD2019.meta 5.ex23&Me) | CLR | -3.54E-10 | 0.955 | 2.20E-10 | 0.955 | -7.29E-11 | 0.955 | -5.65E-10 | 0.691 | -7.13E-10 | 0.563 |
|  | CMS | -1.95E-09 | 0.763 | -2.19E-10 | 0.955 | -1.21E-10 | 0.955 | -1.22E-10 | 0.955 | -1.28E-10 | 0.955 |
|  | iHS | -6.15E-09 | 0.558 | -4.77E-09 | 0.426 | -4.64E-09 | 0.285 | -4.61E-09 | 0.183 | -4.31E-09 | 0.163 |
|  | LA | -2.98E-09 | 0.111 | -1.78E-09 | 0.263 | -1.10E-09 | 0.426 | -1.23E-09 | 0.340 | -1.06E-09 | 0.422 |
|  | XP-EHH | -1.12E-08 | **0.035** | -7.78E-09 | **0.043** | -6.77E-09 | **0.043** | -5.89E-09 | **0.043** | -4.86E-09 | 0.062 |

**Figure 2.**
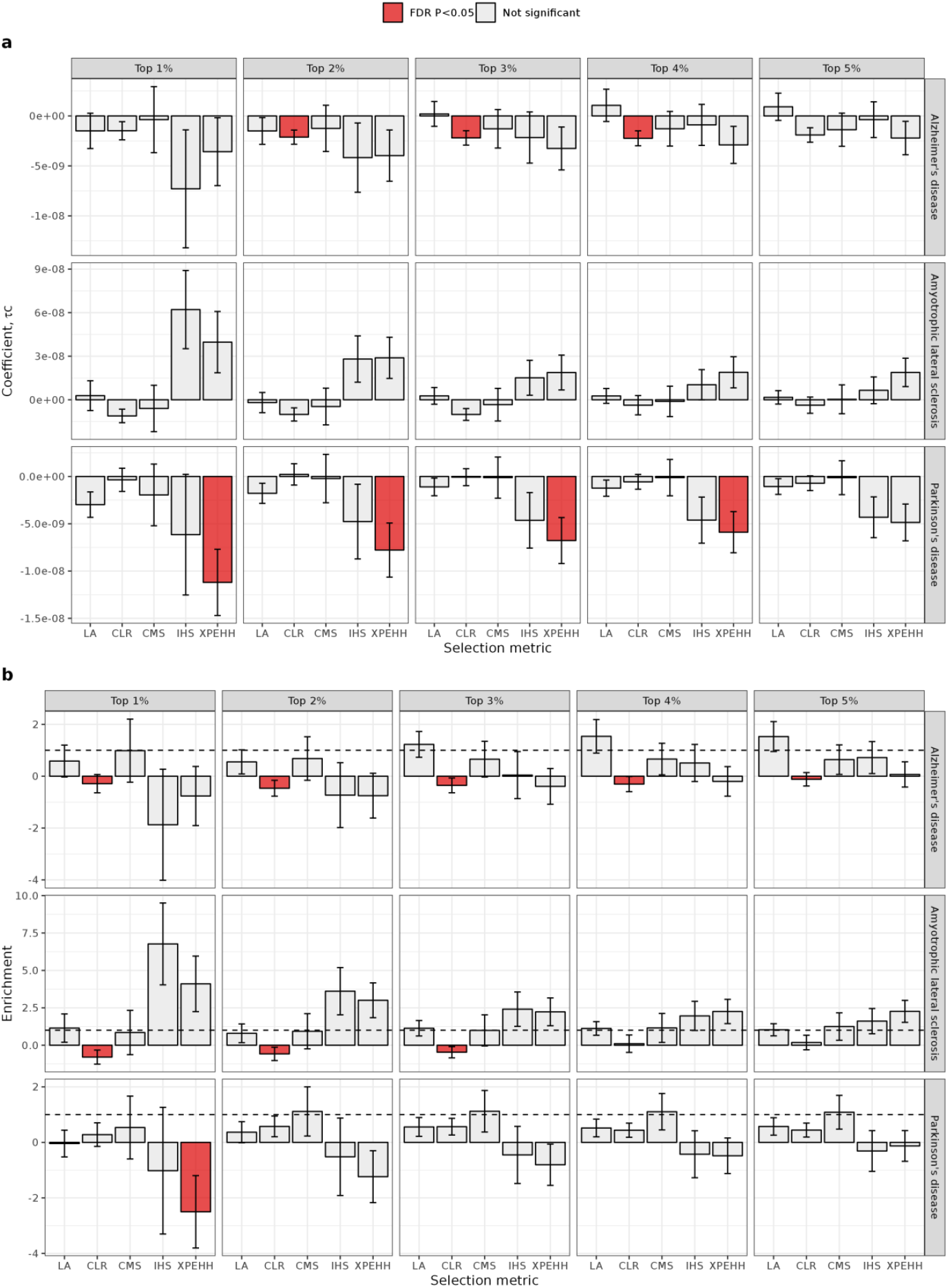
Stratified-linkage disequilibrium score regression analysis across three neurodegenerative disorders comparing SNP-based signatures of Neanderthal introgression and natural selection, stratified across top genome-wide percentiles (1–5%) of annotated SNPs. **(a)The regression coefficient** (*τ*_c_) represents the contribution of the annotation to trait SNP-heritability (*h*^*2*^_SNP_), controlling for all other annotations within the baseline model. The error bars represent 95% confidence intervals. Positive values represent enrichment of trait *h*^*2*^_SNP_ within that annotation. Negative values represent depletion of trait *h*^*2*^_SNP_ within that annotation compared to other annotations within the baseline model. Bars filled with red shading represent significant FDR-corrected two-sided p-values of the regression coefficient (FDR P < 0.05). **(b) Enrichment estimates** from LD score regression analysis. Enrichment values between 0 and 1 indicate a depletion of *h*^*2*^_SNP_ in an annotation category (i.e. showing a lower contribution than expected for a given number of SNPs). Enrichment values larger than 1 indicate an enrichment of *h*^*2*^_SNP_ in an annotation category. The error bars represent 95% confidence intervals. Bars filled with red shading represent significant FDR-corrected two-sided p-values of the regression coefficient (FDR P<0.05). Neanderthal introgression metric (LA) indicates posterior probability of Neanderthal admixture. Positive selection metrics are composite likelihood ratio statistic (CLR), composite of multiple scores (CMS), integrated haplotype score (iHS), cross-population extended haplotype homozygosity (XP-EHH). Genome-wide association studies (GWAS) include AD2019 (Alzheimer’s disease, Jansen *et al*. 2019)^23^, ALS2021.EUR (European cases only from amyotrophic lateral sclerosis GWAS, van Rheenen *et al*. 2021)^24^ and PD2019.met5.ex23&Me (Nalls *et al*. 2019, excluding 23&Me participants).^25^

After correction for multiple testing, we found no evidence for enrichment in Alzheimer’s disease, ALS or Parkinson’s disease *h*^*2*^_SNP_ in alleles most subjected to Neanderthal introgression, as annotated by LA across the top five percentiles (**Figure 2**). This was the case when using both the *τ*_c_ taking into other annotations in the baseline model (**Figure 2a**) and enrichment estimate (**Figure 2b**).

A significant depletion of Parkinson’s disease *h*^*2*^_SNP_ was also observed in SNPs under positive selection, as defined by the top 1 to 4% XP-EHH metric (*τ*_c_:−1.120×10^−8^, −7.780×10^−9^, −6.770×10^−9^, −5.890×10^−9^; FDR P = 0.035, 0.043, 0.043, 0.043 respectively) (**Table 1, Figure 2a**). Importantly, the baseline-LD model controls for metrics of background selection, suggesting that SNPs under positive selection, but under weak or no background selection, are more likely depleted for association with Parkinson’s disease compared to other annotations in the baseline model (**Supplementary Table 4**). However, it should be noted that the *τ*_c_ values are relatively small. Furthermore, given the enrichment estimates for these metrics were negative with larger standard errors towards the top centiles reflecting their lower SNP coverage (0.009% at the top 1% metric annotated by XP-EHH) (**Figure 2b, Supplementary Table 3**), the interpretation of a negative *τ*_c_ here should be interpreted with caution. We then interrogated the contribution of the baseline-LD annotation for background selection for *h*^*2*^_SNP_ in models including each of our natural selection metrics. For each of the five natural selection metrics across all five centiles used, we found that the background selection metric used in the baseline-LD model was not enriched (*τ*_c_ p < 0.05) for Parkinson’s disease *h*^*2*^_SNP_ over and above the enrichment of any of the natural selection metrics and the other 96 baseline-LD annotations (**Supplementary Table 4, Figure 2a**). Therefore, these findings suggest that there is unlikely to be an enrichment of positively-selected SNPs in the genetic architecture of Parkinson’s disease.

A significant depletion of Alzheimer’s disease *h*^*2*^_SNP_ was found using the top 2, 3 and 4% CLR statistic (*τ*_c_ −2.12×10^−9^, −2.20×10^−9^, −2.24×10^−9^; FDR P = 0.023, 0.023, 0.023 respectively). However, there was no nominal depletion across the top 1% of this positive selection metric and given that the corresponding enrichment values are negative, this finding is not conclusive in suggesting that SNPs under positive selection are depleted for association with Alzheimer’s disease (**Figure 2**). Given that there was no evidence for other metrics of natural selection to be enriched for Alzheimer’s disease *h*^*2*^_SNP_, it is also unlikely that positive selection contributed to the evolution of common variants contributing towards the risk of this disease.

After correction for multiple testing, no significant enrichment or depletion of ALS *h*^*2*^_SNP_ were observed in metrics of positive selection, suggesting that positive selection did not play a role in the common genetic risk of ALS.

## Discussion

Humans are particularly vulnerable to neurodegeneration.^1,2^ For example, in ALS, the most frequent neurodegenerative disease of mid-life, motor neurone degeneration results in disruption of multiple motor functions that are key to survival, and therefore of importance for evolutionary adaptation.^40^ Any genetic variation predisposing to motor neurone degeneration might therefore be expected to be under major negative selection pressures. Thus, several possible evolutionary explanations may exist for common alleles contributing to neurodegenerative disease risk to persist in the population despite their deleterious effects.

First, any genetic variation predisposing to disease could have a corresponding benefit and therefore be positively selected. This is a mechanism by which the persistence of Neanderthal-derived sequences in the modern Eurasian human genome has been explained.^14^ For example, a Neanderthal haplotype associated with protection against severe forms of SARS-CoV-2 infection (and other RNA viruses) is also linked to Alzheimer’s disease risk.^41,42,43^ Alternatively, deleterious variants in LD with an advantageous allele may have hitchhiked during positive selection, rising in frequency in the population.^44^ This would be consistent with the proposed Northern founders of the p.91D>A *SOD1* variant in ALS^45^ and the pathogenic *C9orf72* hexanucleotide repeat expansion associated with ALS and frontotemporal dementia^46,47^ Secondly, because neurodegenerative disorders are diseases of ageing, any genetic susceptibility might act after child-rearing years, and therefore outside the age window in which negative selection pressure could have an impact on allele frequency.^48^ In this scenario, the negative effect of the genetic variant is mitigated by the timing of neurodegeneration. Thirdly, neurodegenerative diseases might result from multiple rare variants, each unique in the affected person, and therefore be too rare for selection to have a significant impact.^49^ With the increased availability of high-depth next generation sequencing, many rare variants have now been found to be associated with ALS^24,50,51^, Parkinson’s disease^25,52^ and Alzheimer’s disease^53,54^, but common variants also contribute to risk. To address these hypotheses of how natural selection may have led to the persistence of common genetic risk variants of neurodegenerative disorders, we used stratified-LDSC to test the relationship between neurodegeneration-related SNPs and SNP-based signatures of natural selection.

We found no significant enrichment of Alzheimer’s disease, ALS or Parkinson’s disease *h*^*2*^_SNP_ associated with Neanderthal admixture. This observation is consistent with recent studies using UK Biobank data that showed introgressed variants were depleted for contribution to the heritability of most complex traits or at least not enriched.^21,22^ These studies did not specifically focus on neurodegenerative diseases. Our findings suggest that Neanderthal admixture is unlikely to have maintained the common genetic variant architecture of these neurodegenerative diseases in modern humans.

Positive selection is seen as an evolutionary mechanism for adaptation to a new environment.^39^ In this situation, the beneficial allele sweeps to high frequencies and towards fixation (defined as 100% frequency) together with other variants on the same haplotype, reducing the population genetic diversity in a selective sweep.^39^ SNPs under positive selection were found to be depleted for association with Parkinson’s disease *h*^*2*^_SNP_ compared to other annotations in the baseline model (*τ*_c_) but the simultaneous presence of a negative enrichment estimate suggests that there is at least no evidence of enrichment for positive selection in the heritability of these diseases. This suggests that positive selection does not maintain common risk alleles in Parkinson’s disease. Heritability needs to be considered using this approach to estimate the contribution of natural selection to disease risk. The negative enrichment estimates may also, in part, stem from the relatively low heritability of Parkinson’s disease compared to other neurodegenerative and psychiatric diseases (0.22 according to estimates from genome-wide meta-analysis)^25^, making it more difficult to definitely interpret the depletion in *τ*_c_ as a true depletion in Parkinson’s disease *h*^*2*^_SNP_. Therefore, given the relatively low heritability of Parkinson’s disease, a strong environmental component is implicated in the contribution to disease. This would mean that natural selection would have a limited role in Parkinson’s disease risk. The lack of enrichment of alleles that have been positively selected through Neanderthal admixture or other selective sweeps implicate a strong gene-environment interaction. Indeed, gene-environment interaction frameworks have previously been proposed for the persistence of deleterious common variants in the population gene pool.^55^

In Alzheimer’s disease, this depletion of positively selected SNPs was even less clear, indicating that positive selection is unlikely to be associated with persistence of these common disease-associated variants.

For ALS, we saw no evidence for the enrichment of positive selection in contributing to its trait heritability. This relationship with positive selection is intriguing, providing support for an increased common allele frequency despite the selective removal of alleles that are deleterious, perhaps in combination with mitigation from the timing of disease onset. This is consistent with a previous study that uses UK Biobank data that found positive selection, as tagged by CMS, is not enriched for heritability of neurological traits.^56^ In addition, these findings showing the lack of enrichment from positive selection imply that the genetic architecture of Parkinson’s and Alzheimer’s diseases, and ALS, may stem from multiple, rare, pleiotropic variants rather than polygenic variants with small effects.

This study has several limitations. Firstly, our analyses only studied individuals of European ancestry while Neanderthal introgression also occurred in non-European populations, where the extent of introgression may have differed and thus affects the sensitivity of the analysis. Secondly, the *h*^*2*^_SNP_ estimates using stratified-LDSC analysis are limited by the quality of LD information underpinning the heritability calculations^30^ and the sample size of the GWAS, although we attempted to use only well-powered studies. LDSC also does not take into account the major histocompatibility complex region while Neanderthal alleles have been shown to play a role in modern immune function, thus missing those variants with pleiotropic effects in the nervous system.^16^ We are also excluding rarer Neanderthal introgressed and positively-selected variants by using a MAF cut-off and thus are not able to conclude on the heritability contribution of these variants with lower population frequency. Thirdly, each metric of natural selection and Neanderthal admixture has its own strengths and shortcomings and our analysis is limited by these measures.^39^ However, we attempted to overcome this issue by using a range of SNP-based signatures.^26^ We also note that the negative enrichment estimates are difficult to interpret in this context, possibly due to the SNP coverage of the annotations, which we have tried to compensate for by using higher percentiles of each selection metric. Lastly, this analysis did not take into account structural variants, short tandem repeats or other repetitive genomic elements that may have been acquired through positive selection. A recent study showed a number of repeat elements have risen *ab initio* in *Homo sapiens* which may have implications for positive selection in disease given that these are the most mutable regions of the genome.^57^

In this analysis, we quantified the contribution of positive selection and Neanderthal introgression to the heritability of Alzheimer’s disease, ALS and Parkinson’s disease using stratified-LDSC. We found no significant enrichment of Neanderthal introgression in the SNP-heritability of these neurodegenerative diseases. We also found that genomic regions with positive selection showed no evidence for the contribution in Alzheimer’s disease, ALS and Parkinson’s disease SNP-heritability. This suggests that positive selection does not maintain common risk variants for these disorders in the population even after controlling for background selection. These findings provide further insight into the evolution of the genetic architecture of these disorders.

## Abbreviations

(SNPs): single nucleotide polymorphisms
(ALS): amyotrophic lateral sclerosis
(GWAS): genome-wide association studies
(LDSC): linkage disequilibrium score regression
(LD): linkage disequilibrium
(FDR): False discovery rate

## Contributorship statement

ZC, MRT, MR and AAC conceived and designed the study. ZC, RHR and SAGT critically analysed and interpreted the data. RHR set up and ran the stratified-LDSC analysis pipeline. AFP provided metrics for natural selection. WvR provided summary statistics from the ALS GWAS used. ZC wrote the first draft of the manuscript. JAH, HH, MJO, MRT, MR and AAC supervised the study. KL, AS, EKG, IF and ARJ contributed to the analyses. WR, PC, AC, PJS, KEM, JHV, LHvdV, CES, JFP, and VS contributed to GWAS data used in this study. All authors approved the manuscript and contributed to generation of the submitted draft.

## Acknowledgements

Z.C. is funded by a Leonard Wolfson Foundation clinical research fellowship in neurodegeneration. I.F. receives salary support from the National Institute for Health Research (NIHR) Dementia Biomedical Research Unit at South London and Maudsley NHS Foundation Trust and King’s College London. I.F. is also funded by the Motor Neurone Disease Association (Grant number 905– 793, 6058). W.R. is supported through the E. von Behring Chair for Neuromuscular and Neurodegenerative Disorders, the Laevers Fund for ALS Research, the ALS Liga België, the fund ‘Een Hart voor ALS’ and the fund ‘Opening the Future’. MRT is supported by the Motor Neurone Disease Association. AAC is an NIHR Senior Investigator (NIHR202421). This is an EU Joint Programme - Neurodegenerative Disease Research (JPND) project. The project is supported through the following funding organisations under the aegis of JPND - www.jpnd.eu (United Kingdom, Medical Research Council (MR/L501529/1; MR/R024804/1) and Economic and Social Research Council (ES/L008238/1)) and through the Motor Neurone Disease Association, My Name’5 Doddie Foundation, and Alan Davidson Foundation. This study represents independent research part funded by the National Institute for Health Research (NIHR) Biomedical Research Centre at South London and Maudsley NHS Foundation Trust and King’s College London. The work leading up to this publication was funded by the European Community’s Health Seventh Framework Programme (FP7/2007–2013; grant agreement number 259867 and 340429). The views expressed are those of the authors and not necessarily those of the NHS, the NIHR or the Department of Health.

## Competing interests

AAC reports consultancies or advisory boards for Amylyx, Apellis, Biogen, Brainstorm, Cytokinetics, GenieUs, GSK, Lilly, Mitsubishi Tanabe Pharma, Novartis, OrionPharma, Quralis, and Wave Pharmaceuticals.**Patient consent for publication:** Not applicable.

## Ethics approval

This study does not involve human participants.

## Data availability statement

All data relevant to the study are included in the article or uploaded as supplementary information. Code is available from: https://github.com/RHReynolds/als-neanderthal-analysis.

## Notes

### Summary of Updates

In this revised version, we have updated the results with the 1st to 5th centiles for all annotations to improve SNP-coverage in the LDSC regression analysis to reduce the chance of model misspecification. We have also refrained from commenting on nominally enriched p-values.

